# The relative densities of cell cytoplasm, nucleoplasm, and nucleoli are robustly conserved during cell cycle and drug perturbations

**DOI:** 10.1101/2020.04.14.040774

**Authors:** Kyoohyun Kim, Jochen Guck

## Abstract

The cell nucleus is a compartment in which essential processes such as gene transcription and DNA replication occur. While the large amount of chromatin confined in the finite nuclear space could install the picture of a particularly dense organelle surrounded by less dense cytoplasm, recent studies have begun to report the opposite. However, the generality of this newly emerging, opposite picture has so far not been tested. Here, we used combined optical diffraction tomography (ODT) and epi-fluorescence microscopy to systematically quantify the mass densities of cytoplasm, nucleoplasm, and nucleoli of human cell lines, challenged by various perturbations. We found that the nucleoplasm maintains a lower mass density than cytoplasm during cell cycle progression by scaling its volume to match the increase of dry mass during cell growth. At the same time, nucleoli exhibited a significantly higher mass density than the cytoplasm. Moreover, actin and microtubule depolymerization and changing chromatin condensation altered volume, shape, and dry mass of those compartments, while the relative distribution of mass densities was generally unchanged. Our findings suggest that the relative mass densities across membrane-bound and membraneless compartments are robustly conserved, likely by different as of yet unknown mechanisms, which hints at an underlying functional relevance. This surprising robustness of mass densities contributes to an increasing recognition of the importance of physico-chemical properties in determining cellular characteristics and compartments.

## Introduction

The physical properties of cells and their membrane-bound and membraneless compartments are increasingly becoming the focus of current cell and developmental biological research (1, 2). The cell nucleus is a prominent example of a membrane-bound compartment that maintains physical and biochemical conditions distinct from the surrounding cytoplasm. Alterations of nuclear physical properties are associated with various diseases (3). Nuclei also harbor nucleoli within which ribosomal subunits are synthesized and assembled for protein translation. Nucleoli are prime examples of membraneless compartments being dynamically maintained by liquid-liquid phase separation (4, 5). Due to high levels of metabolic activity, it might be conceivable that the nucleolus is less dense than the surrounding nucleoplasm, while a previous study reported that nucleoli consist of a fibrillar region with higher density than the nucleoplasm, surrounded by low-density, sponge-like granular components (6). And, because of the large amount of DNA, histones, and various other types of proteins tightly packed into the finite space of the nucleus, the nucleus has been perceived as a particularly dense organelle compared to the less dense cytoplasm (7–9).

Recent studies, however, have started to paint a different picture. Using quantitative phase imaging and optical diffraction tomography, the cell nucleus was reported to have a lower refractive index (RI) than cytoplasm (10–14). Since the RI in most biological materials is linearly proportional to their mass density (15, 16), the results indicated that the cell nucleus also has lower mass density than cytoplasm. This finding is supported by other approaches measuring RI, including surface plasmon resonance microscopy (17), transport-of-intensity microscopy (18), and orientation-independent differential interference microscopy (19), but also other indirect approaches to quantify mass density such as Raman microscopy (20). For a generalization of this finding, though, important questions still remain open. In particular, mass density distributions could change during the cell cycle when the genetic material and protein machinery in the nucleus is being duplicated, and more mass added to the nucleus during S/G2 phase. Nuclear volume, and thus density, could depend on the tensional state of the nucleus via the cytoskeleton (21, 22). Or, chromatin density might be a direct function of epigenetic markers, such as methylation or acetylation of histone proteins, controlling chromatin condensation (23). Perturbing these aspects permits testing the robustness of the phenomenon.

Here, we systematically investigated the relative densities of nucleoplasm, cytoplasm, and nucleoli in two different adherent mammalian cell types (HeLa-FUCCI, RPE-FUCCI) by quantifying their 3D RI distributions using combined optical diffraction tomography (ODT) and epi-fluorescence microscopy. By correlating the RI tomograms of cells with epi-fluorescence images of their nuclei stained with the cell-cycle dependent FUCCI (fluorescent ubiquitination-based cell cycle indicator) fluorescence markers, we quantitatively characterized the RI, mass density, dry mass, and volume of nucleoplasm, nucleoli, and cytoplasm for the different cell-cycle phases. We observed that — throughout the cell cycle — nucleoplasm has a lower RI and mass density than cytoplasm. This does not change as the volume of the nucleoplasm and cytoplasm scales with the increase of dry mass as the cell grows and duplicates its genetic material. In contrast, nucleoli inside the nucleus exhibit significantly higher mass density than both nucleoplasm and cytoplasm. This relative mass density distribution between compartments was robust against considerable drug-induced perturbations such as depolymerization of actin and microtubules as well as chromatin condensation and decondensation, even though the shape, density, volume and dry mass of each compartment of course changed. The robustness of this surprising finding suggests that there might be an underlying functional relevance and that there are as of yet unknown active, or passive, mechanisms that maintain relative mass densities across both membrane-bound and membraneless compartments.

## Materials and Methods

### Cell culture and preparation

The stable HeLa-FUCCI and RPE-FUCCI cell lines were kindly provided by the lab of Frank Buchholz (TU Dresden). FUCCI is a fluorescent ubiquitination-based cell cycle indicator (24). The stable HeLa cell line transfected with GFP:NIFK, which labels nucleolar protein interacting with the FHA domain of pKI-67, was kindly provided by the lab of Anthony Hyman (Max Planck Institute of Molecular Cell Biology and Genetics, Dresden). All cell lines were cultured under standard conditions at 37°C, 5% CO_2_. HeLa cells were cultured in standard DMEM, high glucose with GlutaMax medium (61965-026, Thermo Fisher Scientific, Inc.), and RPE cells were cultured in DMEM/F12, high glucose with GlutaMax medium (31331-028, Thermo Fisher Scientific, Inc.). All culture media were supplemented with 10% FBS and 1% penicillin-streptomycin. The cells were subcultured in a glass-bottom petri dish (FluoroDish, World Precision Instruments GmbH) one day prior to the measurement, and the culture media were exchanged to CO_2_ independent medium (18045-088, Thermo Fisher Scientific, Inc.) prior to imaging. HeLa-GFP:NIFK cells were stained with Hoechst (1:1,000 dilution) for nuclei staining and washed with fresh CO_2_-independent medium prior to imaging.

### Drug treatments

In order to investigate the role of the cytoskeleton on maintaining the RI of the nucleus and cytoplasm, cells were treated with 1 μM cytochalasin D (cytoD) for 30 minutes or 5 μM nocodazole (noco) for 1 hour before imaging to depolymerize actin and microtubules, respectively. To decondense chromatin, cells were incubated with 300 nM trichostatin A (TSA), a histone deacetylase inhibitor, in cell culture medium for 9 hours. To condense chromatin, cells were incubated with 100 μM anacardic acid (ANA), a histone acetyltransferase inhibitor, in cell culture medium for 1 hour.

### Optical setup

The optical setup consisted of a combined ODT and epi-fluorescence microscope (Fig. 1) as described previously (12). Briefly, ODT employs Mach-Zehnder interferometry in order to measure the 3D RI distribution of cells (Fig. 1 *a*). A laser beam (*λ* = 532 nm, frequency-doubled Nd-YAG laser, Torus, Laser Quantum Inc.) was coupled into an optical fiber and divided into two beams by a 2 × 2 single-mode fiber-optic coupler.

**Figure 1.**
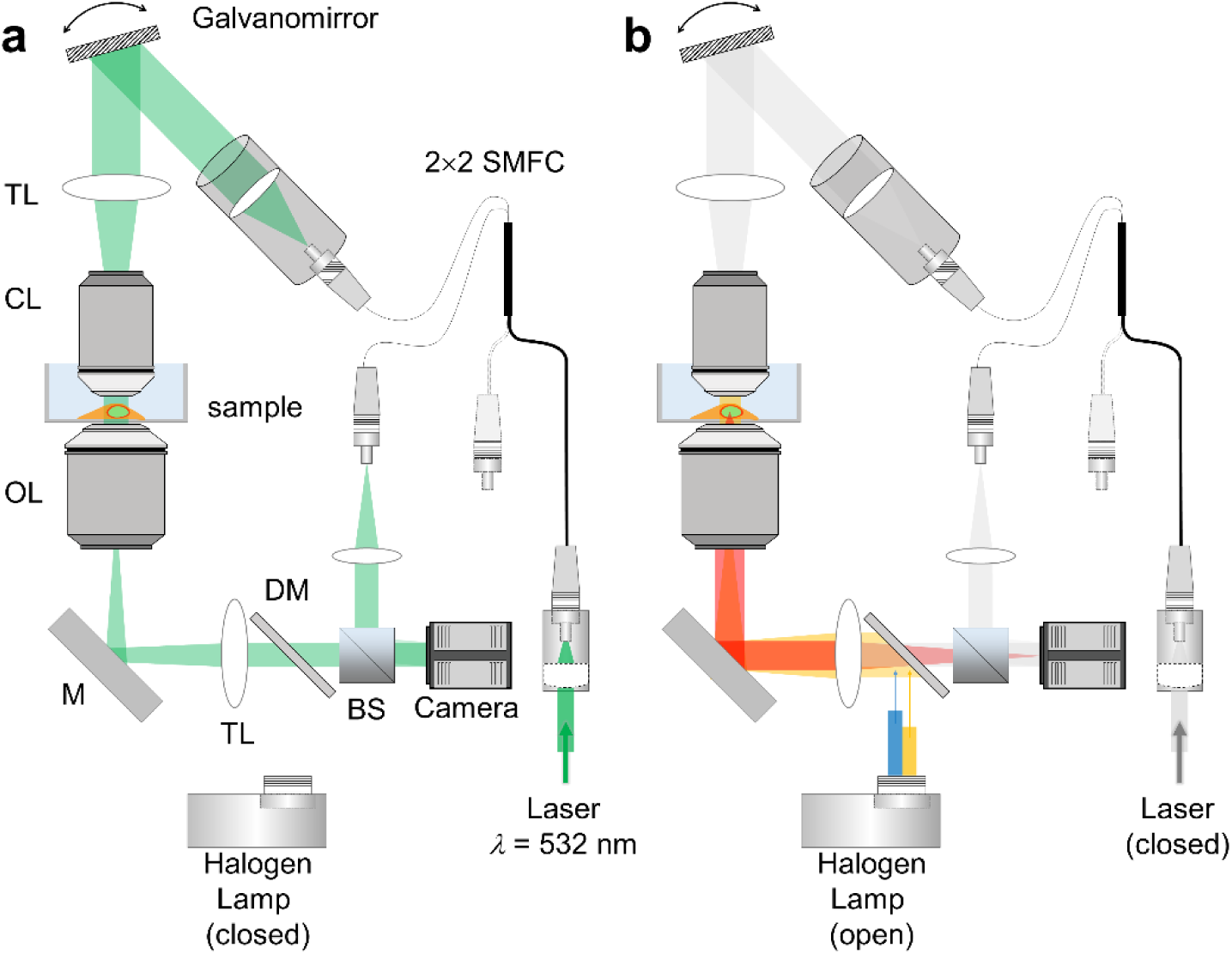
Experimental setup. (a) The optical setup for optical diffraction tomography (ODT). SMFC, single-mode fiber coupler; TL, tube lens; CL, condenser lens; OL, objective lens; M, mirror; DM, dichroic mirror; BS, beam splitter. (b) The same optical setup when used for epi-fluorescence microscopy.

One beam was used as a reference beam and the other beam illuminated the cells on the stage of a custom-made inverted microscope through a tube lens (*f* = 175 mm) and a water-dipping objective lens (NA = 1.0, 40×, Carl Zeiss AG). To reconstruct a 3D RI tomogram of cells, the samples were illuminated from 150 different incident angles scanned by a dual-axis galvano-mirror (GVS012/M, Thorlabs Inc.). The beam diffracted by the sample was collected by a high numerical-aperture objective lens (NA = 1.2, 63×, water immersion, Carl Zeiss AG) and a tube lens (*f* = 200 mm). The total magnification was set to be 57×. The diffracted beam interfered with the reference beam at an image plane, and generated a spatially modulated hologram, which was recorded with a CCD camera (FL3-U3-13Y3M-C, FLIR Systems, Inc.). In order to maintain the temperature of the culture medium in the glass-bottom petri dish, both objective lenses were heated at 37°C by resistive foil heaters (Thorlabs Inc.).

Epi-fluorescence imaging was performed using the same optical setup (Fig. 1 *b*). In order to excite fluorescence probes in cells, an incoherent beam from a halogen lamp (DC-950, Dolan-Jenner Industries Inc.) was coupled into the same beam path using a three-channel dichroic mirror (FF409/493/596-Di01-25×36, Semrock Inc.). The excitation wavelength was selected by alternating bandpass filters corresponding to Hoechst, monomeric Azami-Green 1 (mAG1) and monomeric Kusabira-Orange 2 (mKO2).

### Tomogram reconstruction and quantitative analysis

The complex optical fields of light scattered by the samples were retrieved from the recorded holograms by applying a Fourier transform-based field retrieval algorithm (25). The 3D RI distribution of the samples was reconstructed from the retrieved complex optical fields via the Fourier diffraction theorem (26, 27). A more detailed description of tomogram reconstruction can be found elsewhere (28, 29). The spatial resolution of ODT is 121 nm (lateral) and 444 nm (axial), which was determined by the NAs of the objective lens and condenser lens (30). The precision of the ODT was characterized by measuring the standard error of RI tomograms of the same FOV with repetitive measurements (Supplementary Figure 1). We measured a time series of 10 RI tomograms of silica beads immersed in 0.7 M sucrose solution (*n* = 1.3665 at *λ* = 532 nm) in the same FOV with the tomogram acquisition rate of 1 Hz. We calculated the standard error of the time series of 10 RI tomograms, and averaged the standard error within individual silica bead. The averaged standard error of RI was calculated as 4.15 × 10^−5^, which can be converted to the error of the absolute mass density in cells as 0.22 mg/ml.

From the reconstructed tomograms (Fig. 2 *a*), Otsu’s thresholding method (31) was used to segment the regions occupied by cells from background, and the watershed algorithm to identify individual cells from segmented RI tomograms. Then, epi-fluorescence images of nuclei stained with the FUCCI marker (Fig. 2 *b*) were correlated with the RI tomograms in order to segment nuclei inside cells. We also segmented the periphery of nuclei, *i*.*e*., perinuclear cytoplasm where the endoplasmic reticulum and Golgi apparatus are dominant, by dilating the binary nuclei masks by 2 μm thickness. Inside the 3D RI distribution of nuclei, regions having higher RI values than surrounding nucleoplasm were segmented by Otsu’s thresholding method and identified as nucleoli. We confirmed that nucleoli had higher RI than nucleoplasm by correlating the RI tomograms of HeLa cells with epi-fluorescence images where nuclei and nucleoli were stained with Hoechst and GFP, respectively. The number of voxels in the region of nucleoli segmented by the present RI-based method had 99.5% true positive rate (sensitivity) compared to the epi-fluorescence images of nucleoli, and the nucleoplasm could be segmented with 88.9% sensitivity (see Supplementary Figure 2). Finally, we quantified the 3D RI distribution of cytoplasm (*n*_c_), perinuclear cytoplasm (*n*_pc_), nucleoplasm (*n*_np_), and nucleoli (*n*_nl_) separately from the segmented tomogram (Fig. 2 *c*-*f*).

**Figure 2.**
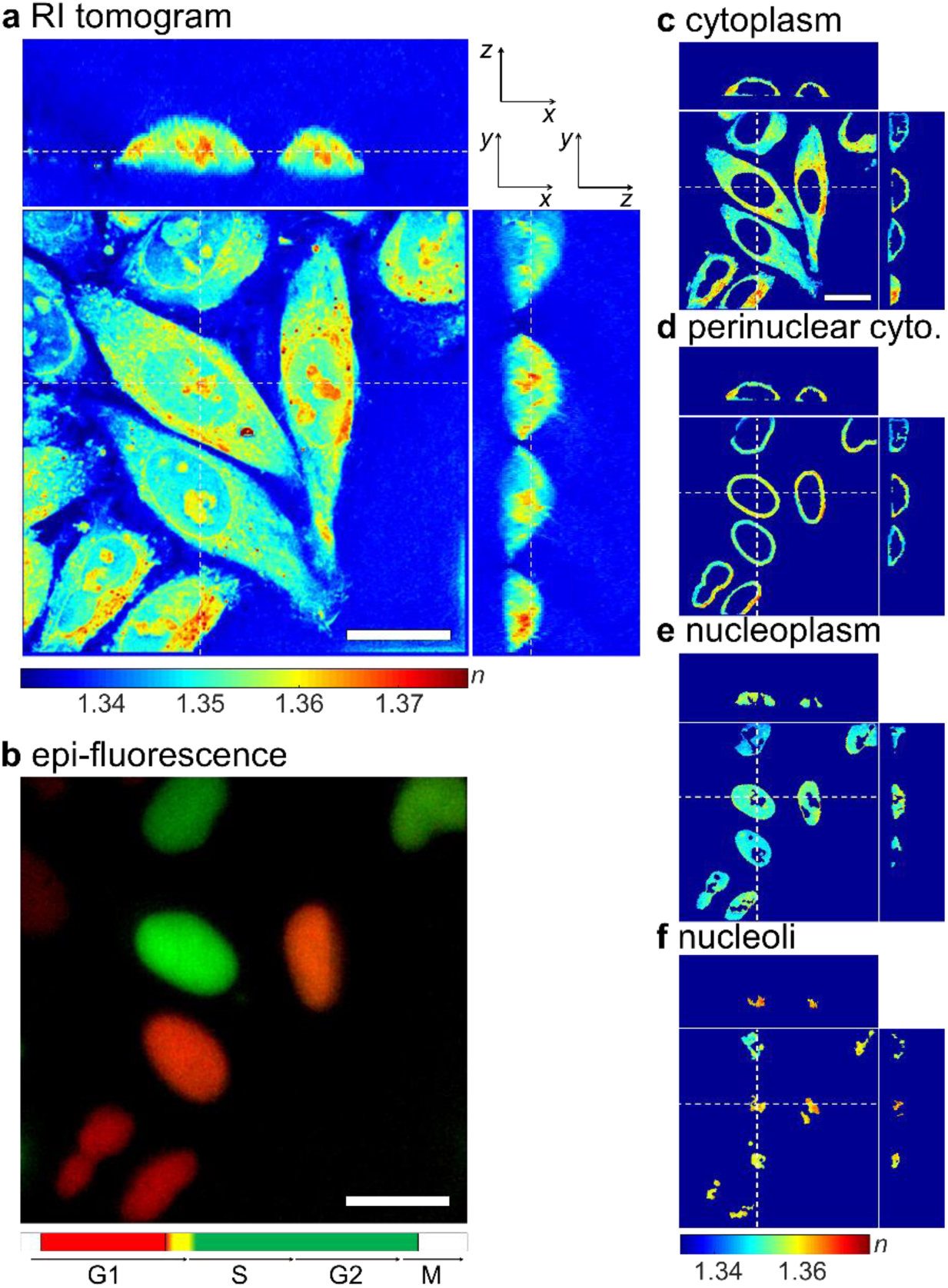
Segmentation of cellular and nuclear compartments from an RI tomogram and epi-fluorescence map. (a) The cross-sectional slices of an RI tomogram of HeLa-FUCCI cells in the *x*-*y, x*-*z*, and *y*-*z* planes. The color scale quantifies the refractive index, *n*. (b) The epi-fluorescence map of the same field-of-view as in (a). Red color identifies cells in the G1 phase of the cell cycle, yellow in early S, and green in S/G2. (c-f) The RI maps segmented for (c) cytoplasm, (d) perinuclear cytoplasm, (e) nucleoplasm, and (f) nucleoli. Dashed lines indicate the corresponding *x*-*z* and *y*-*z* planes. Scale bars are 20 µm.

The mass density of each compartment was directly calculated from the mean RI value, since the RI value in biological samples, *n*(*x,y,z*), is linearly proportional to the mass density of the material, *ρ*(*x,y,z*), as *n*(*x,y,z*) = *n*_m_ + *αρ*(*x,y,z*), where *n*_m_ is the RI value of the surrounding medium and *α* is an RI increment (*dn/dc*). The RI of the medium was measured using an Abbe refractometer (2WAJ, Arcarda GmbH) to be *n*_m_ = 1.3370 ± 0.00025 at *λ* = 532 nm. The RI increment used was *α* = 0.190 mL/g for protein and nucleic acid (32, 33). In addition, the volume of the compartments was extracted by counting the number of voxels in the corresponding binary mask, and the dry mass of the compartments was calculated by integrating the mass density inside the corresponding binary mask. The sphericity of the compartments, *Ψ*, was calculated as the ratio between surface area, *A*, and volume of the mask, *V*, as *Ψ* = [π^1/3^(6*V*)^2/3^]/*A*. All tomogram acquisition and data analysis was performed using custom-written MATLAB scripts (R2017b, MathWorks, Inc.), which are available in GitHub (https://github.com/OpticalDiffractionTomography/NucleiAnalysis).

### Statistical Analysis

We tested the normality of the RI distributions and the RI ratios using the Shapiro-Wilk test, which revealed that most distributions were normal. We report means and standard error of means (SEM) for the data passing the normality test, and means when the data did not pass the normality test. Statistical significance between data groups was determined using either the two-tailed Student’s *t*-test, or the two-tailed Mann-Whitney *U*-test when the data did not pass the normality test. The shown asterisks indicate the statistical significance as \**p* < 0.01, \*\**p* < 0.001, and \*\*\**p* < 0.0001, respectively.

## Results

### Relative compartment RI and mass densities are independent of cell cycle and cell types

In order to investigate the effect of the cell cycle on the RI distribution of cytoplasm, nucleoplasm, and nucleoli, we correlated the reconstructed RI tomograms of two adherent cell lines, HeLa-FUCCI and RPE-FUCCI cells, with the respective FUCCI-fluorescence images. We grouped each cell into the G1, early S, and S/G2 phases of the cell cycle (*N* = 557, 505, and 483 for HeLa cells, and *N* = 122, 92, and 128 for RPE cells in the G1, early S, and S/G2 phases, respectively) by the average intensity of their epi-fluorescence images in the mAG1 and mKO2 channels (see Supplementary Figure 3 for details). As shown in the typical examples in Figure 3 *a*-*b* and Figure 3 *d*-*e*, the cell nucleoplasm of both HeLa and RPE cells had lower RI values compared to the cytoplasm in all phases of the cell cycle. Staining with FUCCI dyes did not affect the RI of the compartments as non-labeled HeLa cells also showed similar RI distribution (Supplementary Figure 4).

**Figure 3.**
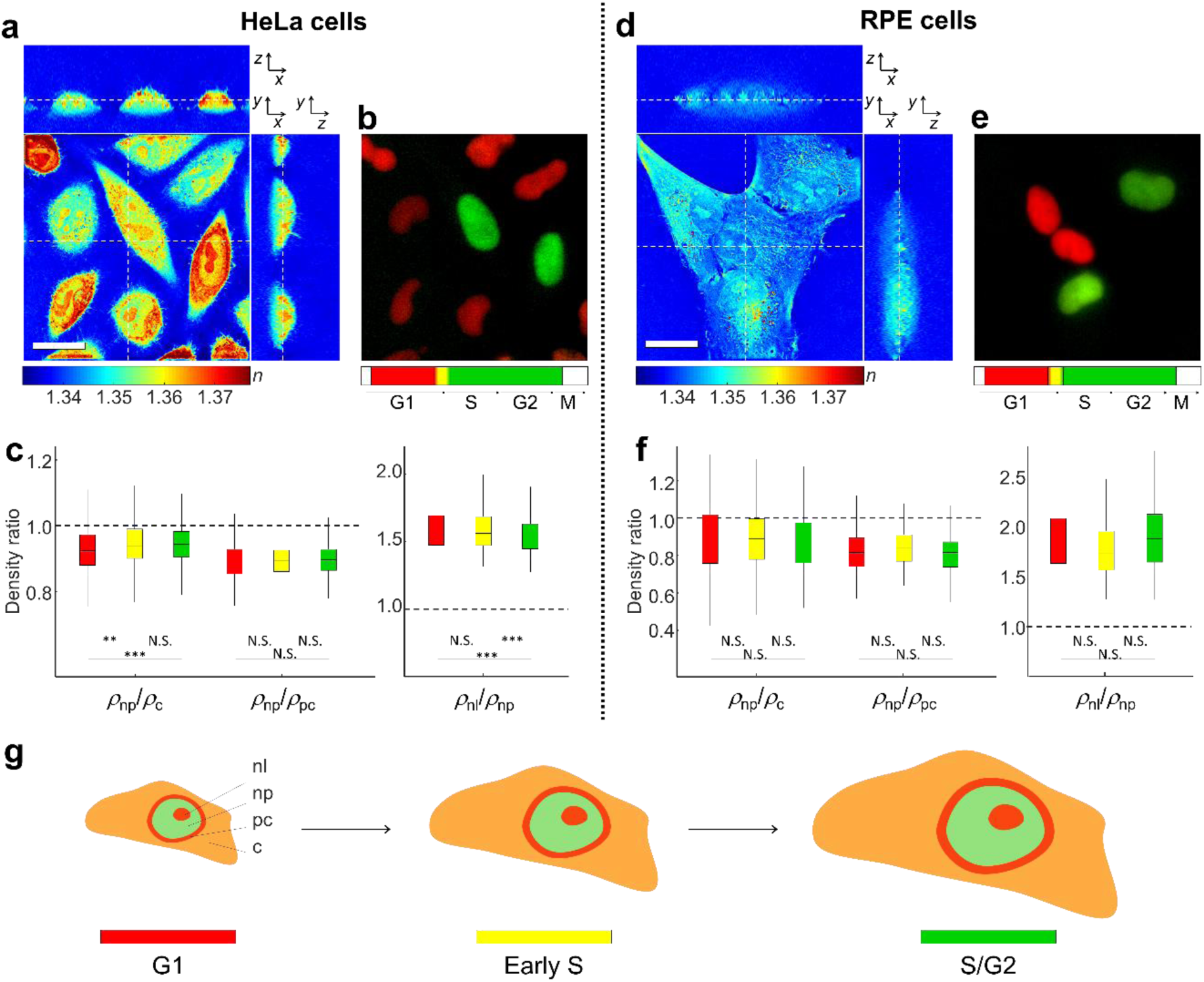
Refractive index (RI) and mass density distributions in HeLa-FUCCI (a-c) and RPE-FUCCI (d-f) cells as function of the cell cycle. (*a, d*) Cross-sectional slices of 3D RI tomograms of cells in the *x*-*y, x*-*z*, and *y*-*z* planes. The dashed lines indicate the corresponding *x*-*z* and *y*-*z* planes. The color scale quantifies the refractive index, *n*. Scale bars are 20 µm. (*b*-*e*) Corresponding FUCCI-fluorescence images of nuclei in different cell cycle phases. Red color identifies cells in the G1 phase of the cell cycle, yellow in early S, and green in S/G2. (*c*-*f*) The ratio of the mass density between nucleoplasm and cytoplasm (*ρ*_np_/*ρ*_c_), nucleoplasm and perinuclear cytoplasm (*ρ*_np_/*ρ*_pc_), and nucleoli and nucleoplasm (*ρ*_nl_/*ρ*_np_). The dashed lines indicate equal mass density between compartments, and the box plots represent the interquartile ranges (IQR) with a line at the median. The colors indicate cell cycle phases as in (*b, e*). The numbers of cells measured are *N* = 557, 505, and 483 for HeLa cells, and *N* = 122, 92, and 128 for RPE cells in the G1, early S, and S/G2 phases, respectively. (*g*) Schematic indicating the four segmented regions (cytoplasm: c, perinuclear cytoplasm: pc, nucleoplasm: np, and nucleoli: nl) and how they scale in size to maintain the same relative mass densities throughout the cell cycle.

For a more quantitative analysis, we calculated the mean RI values, and the mean mass densities, of cytoplasm, perinuclear cytoplasm (as a part of the cytoplasm), nucleoplasm, and nucleoli from the segmented RI tomograms of individual cells (see Methods; Supplementary Figure 5). We found that nucleoplasm of HeLa cells was 7.3 ± 0.6%, 5.8 ± 0.6%, and 5.7 ± 0.6% (mean ± SEM) less dense than cytoplasm at the G1, early S, and S/G2 phases, respectively (Fig. 3 *c*). The absolute mass density differences of 6.0 ± 0.5 mg/ml, 4.9 ± 0.5 mg/ml, and 4.7 ± 0.5 mg/ml were statistically significant (see Supplementary Figure 5 *a, b*). The immediate periphery of the nucleus, *i*.*e*. the perinuclear cytoplasm, where the membrane-rich endoplasmic reticulum and Golgi apparatus are dominant, had even higher RI than other cytoplasmic regions (Supplementary Figure 5 *a, b*). Hence, the relative difference in mass density between cell nucleoplasm and the perinuclear cytoplasm was even more pronounced (10.8 ± 0.5%, 10.6 ± 0.4%, and 10.3 ± 0.4% at the G1, early S, and S/G2 phases, respectively; Fig. 3 *c*).

The RPE cells, as a non-cancerous cell line, showed similar results as nucleoplasm was 10.5 ± 2.9%, 8.7 ± 3.9%, and 13.3 ± 2.7% less dense than cytoplasm, and 18.8 ± 2.0%, 16.4 ± 2.3%, and 19.6 ± 1.6% less dense than the perinuclear cytoplasm at the G1, early S, and S/G2 phase, respectively (Fig. 3 *f*). The absolute mass density differences between nucleoplasm and cytoplasm (4.0 ± 1.2 mg/ml, 3.7 ± 1.8 mg/ml, and 5.2 ± 1.1 mg/ml in the three respective cell cycle phases) was again statistically significant (see Supplementary Figure 5 *c, d*). The lower density of the nucleoplasm compared to the surrounding cytoplasm, thus, seems to be cell-type independent.

In this study, we used the same RI increment, *α* = 0.190 mL/g, to calculate the mass density of each compartment from measured RI, which is valid for protein and nucleic acid (32, 33). However, because the RI increment of phospholipid is lower as *α* = 0.135 – 0.138 ml/g (34, 35), the mass density of such membrane-rich perinuclear cytoplasm may be underestimated and have even higher mass density. Nonetheless nucleoplasm is still less dense than cytoplasm without the perinuclear area under this consideration (the absolute mass density differences of 3.9 ± 1.0 mg/ml, 3.4 ± 1.0 mg/ml, and 3.8 ± 0.9 mg/ml for HeLa cells and 2.8 ± 1.7 mg/ml, 1.8 ± 2.1 mg/ml, and 3.8 ± 1.5 mg/ml for RPE cells in the three respective cell cycle phases, Supplementary Figure 5).

During the cell cycle, cells are growing and duplicating genetic material, which may increase the dry mass, volume, and/or mass density of the nucleus and cytoplasm and could therefore alter our findings. However, both the volume and dry mass of both nucleoplasm and cytoplasm in HeLa and RPE cells increased two-fold during the cell cycle, so that the mass density of both compartments was maintained at about the same level (see Supplementary Figure 6). This result, obtained with two different attached cell lines, is consistent with our previous study in suspended cells (36), and implies that the volume, or the mass density, of the nucleoplasm and cytoplasm is well controlled in response to the increase of cellular content during the cell cycle.

Interestingly, the nucleoli inside the nucleoplasm, where ribosomal subunits are synthesized for protein translation, had significantly higher mean RI and density than the nucleoplasm (58% in HeLa cells, and 92% in RPE cells) and the cytoplasm (48% in HeLa cells and 71% in RPE cells) (Fig. 3 *c, f*; Supplementary Fig. 5 *b, d*). Here, their relative density ratio distribution exhibited the non-normal distribution. Moreover, the averaged mass density of the entire nucleus including nucleoplasm and nucleoli is similar to that of cytoplasm and perinuclear cytoplasm in HeLa cells or even higher than cytoplasm in RPE cells (Supplementary Fig. 7). Recently, several reports have described the nucleolus as a membraneless organelle, which was proposed to assemble by liquid-liquid phase separation (4, 5). Phase separation is a density transition in which two phases — one dilute and one dense phase — stably coexist. Our finding suggest that cells may redistribute the mass within the nucleus by phase separation and formation of the nucleolus and that this generates the relative mass density difference between nucleoplasm and cytoplasm. In addition, our results demonstrate the narrow distributions of the mass density ratio, and difference, between nucleoli and nucleoplasm, which suggests that nucleoli formation and ribosome biogenesis are precisely regulated in these membraneless organelles during the cell cycle.

### Changes due to cytoskeletal perturbations do not abolish relative mass density differences

In the previous section, we have confirmed that nucleoplasm has lower mass density than cytoplasm. We found the same pattern in various cell types, both suspended (13, 36) and attached, and throughout the cell cycle. Apparently, in physiological conditions, the pattern is robustly conserved. But can we identify the origin of this pattern by specific perturbations? There is evidence that the nucleus is under tension from the cytoskeleton, which in turn is anchored to the cell membrane (21, 37). Is it, thus, possible to reduce the volume of the nucleus, enclosing a relatively fixed amount of solid material, by perturbing the tensional balance between cell surface and nucleus in order to increase its density? Cytochalasin D (cytoD) inhibits actin polymerization and destabilizes the actin cortex, while nocodazole (noco) depolymerizes microtubules, and at the same time activates Rho-signaling and increases actomyosin contractility (38). Visual inspection of the reconstructed RI tomograms (Fig. 4 *a*-*c*) showed that HeLa-FUCCI cells treated with cytochalasin D exhibited dramatic morphological changes at the cell boundary (Fig. 4 *b*). Cells treated with nocodazole seemed to be larger and exhibited lower RI values in the cytoplasm (Fig. 4 *c*).

**Figure 4.**
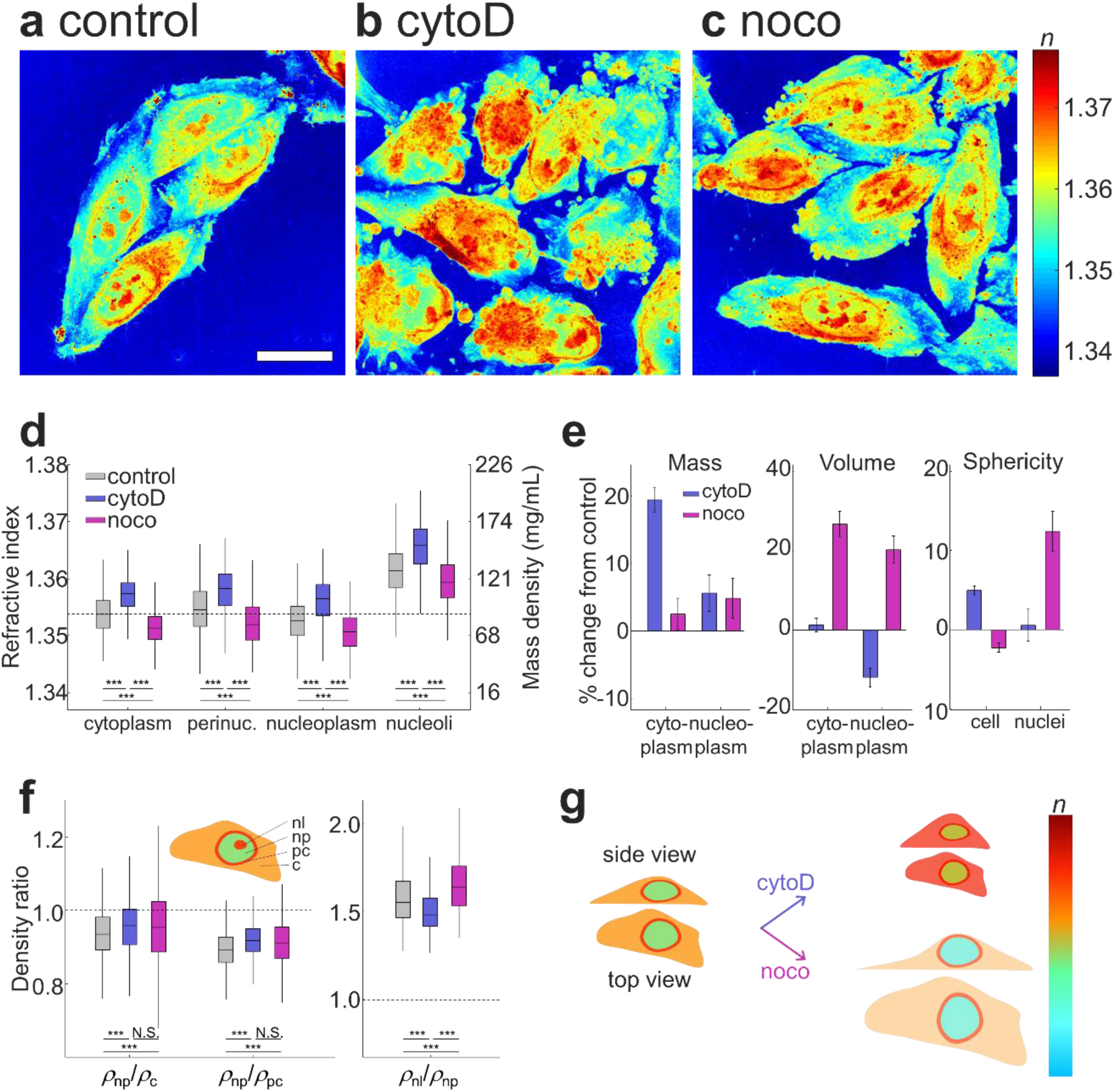
Changes in refractive index distribution of HeLa-FUCCI cells under cytoskeletal perturbation. (a-c) Maximum projection of RI tomograms of HeLa-FUCCI cells in (a) control, (b) cytoD, and (c) noco treatment. The scale bar is 20 µm. (d) Mean RI values of cytoplasm, perinuclear cytoplasm, nucleoplasm, and nucleoli in individual cells under cytoskeletal perturbation. The dashed line indicates the mean RI value of cytoplasm in control cells. (e) Relative changes of the mass, volume, and sphericity of the cytoplasm and nuclei under cytoD and noco treatment compared to control. (f) Mass density ratio between nucleoplasm and cytoplasm (*ρ*_np_/*ρ*_c_), nucleoplasm and perinuclear cytoplasm (*ρ*_np_/*ρ*_pc_), and nucleoli and nucleoplasm (*ρ*_nl_/*ρ*_np_) at the same phases of the cell cycle. The dashed lines indicate equal mass density of the compartments. The numbers of cells measured are *N* = 1,565, 973, and 717 for control, cytoD, and noco treatments, respectively. (g) Schematic indicating the effect of the cytoD and noco treatment on the volume, RI and shape of cells.

Quantitatively, cells treated with cytochalasin D had higher mean RI values in all subcellular compartments compared to controls, while nocodazole-treated cells had lower mean RI values in all compartments (Fig. 4 *d*). Particularly, the mean RI values of cytoplasm and nucleoplasm under the cytoD treatment were 1.3571 ± 0.0002 and 1.3563 ± 0.0002 (mean ± SEM, *N* = 973), and thus higher than controls (1.3538 ± 0.0002 and 1.3528 ± 0.0002 for cytoplasm and nucleoplasm, *N* = 1,565). As shown in Fig. 4 *e*, this increase of mean RI values in the cytoD-treated cells resulted from different physical phenomena. The mean RI value and mass density of nucleoplasm increased as the volume occupied by nucleoplasm decreased 11.9 ± 2.4% compared to controls, while its dry mass only increased by 5.6 ± 2.7%. In contrast, the mean mass density of cytoplasm increased due to the 19.4 ± 1.9% increase of cytoplasmic dry mass while its volume remained the same (1.3 ± 1.7% increases). It is also worth noting that cytoD-treated cells became more spherical as the sphericity of cells increased by 5.0 ± 0.6%, while the nucleus maintained the same sphericity (0.6 ± 2.0% increases). The sphericity of the entire cell can be increased by the detachment from the substrate due to depolymerized actin filaments at the cell boundary, while constant nuclear sphericity during the shrinkage of a nucleus implies that the nuclear volume shrinks uniformly in all directions. These percentage changes are averaged ratios for each cell cycle phase, each of which shows the same change (See Supplementary Figure 8).

Nuclear shrinkage by the cytochalasin D treatment is as expected because cytochalasin D dissociates the actomyosin cytoskeleton, which normally exerts outward tension on the nucleus (39–41). The treatment, then, increased the mass density of nucleoplasm due to the reduced volume at constant dry mass enclosed. At the same time, the mass density of cytoplasm increased due to the increase of its dry mass at about constant volume. Even though there are previous reports that cytochalasin D treatment promotes the synthesis of actin (42) and other proteins (43), it is unclear whether this is the actual mechanism as 30 min of treatment time seems too short to account for 20% mass increase. The actomyosin network can also form a perinuclear actin cap and exert compressive forces on the nucleus in the axial direction, in particular on nuclei in stromal cells (44, 45). In this study, a compressive force on the nucleus by the actomyosin network is probably not relevant as the perinuclear actin cap is absent in HeLa cells (46–48). Taken together, we found that nucleoplasm still had lower mass density than cytoplasm and perinuclear cytoplasm (4.6 ± 0.5% and 8.0 ± 0.3%) (Fig. 4 *f*).

This robustness of the relative mass density between compartments was also observed in cells treated with nocodazole, while the mass RI values of cytoplasm and nucleoplasm decreased as 1.3515 ± 0.0002 and 1.3509 ± 0.0003 (mean ± SEM, *N* = 717), respectively (1.3538 ± 0.0002 and 1.3528 ± 0.0002 for cytoplasm and nucleoplasm in controls). The decreases of mean RI value and the mass density of cytoplasm and nucleoplasm mostly originated from their volume expansion (26.7 ± 3.1% and 20.2 ± 3.4% in cytoplasm and nucleoplasm, respectively) while their dry mass only slightly increased as 2.5 ± 2.3% and 4.8 ± 2.9% (Fig. 4 *e*). The nuclear volume expansion and increased nuclear sphericity in response to the nocodazole treatment can be rationalized based on previous studies, since the microtubules are depolymerized by the nocodazole treatment. Microtubules normally confine the nucleus in the direction perpendicular to the cell culture substrate (41, 49, 50). The effect can be also observed from the increased sphericity of the nucleus increased by 12.4 ± 2.5% while the cell sphericity slightly decreased by 2.3 ± 0.6% (Fig. 4 *e*). The volume increase of the nucleus, at a constant dry mass of the nucleoplasm, then led to a decrease of the nucleoplasm mass density. Interestingly, at the same time, the mass density of cytoplasm also decreased due to the increase of its volume as nocodazole inhibits the normal regulatory volume decrease (49), which in sum again conserved the relative mass density difference between nucleoplasm and cytoplasm. Quantitatively, the nucleoplasm of the nocodazole-treated cells was still 4.3 ± 0.8% less dense than the cytoplasm and 8.5 ± 0.5% less dense than perinuclear cytoplasm (Fig. 4 *f*).

Our finding shows that the relative mass density between nucleoplasm and cytoplasm is still maintained against the perturbation of cytoskeletal integrity, which is graphically summarized in Fig. 4 *g*. We also revealed that the mass density difference across the nuclear membrane of cytochalasin D- and nocodazole-treated cells was 8.8 ± 0.4 mg/ml and 7.1 ± 0.4 mg/ml, which are slightly lower than that of control cells as 10.2 ± 0.3 mg/ml. It may imply that cytoskeletal perturbation may induce a subtle force imbalance between the osmotic pressure gradient across the nuclear membrane and the force exerted by the cytoskeleton (50– 52).

### Perturbing chromatin condensation still does not overturn relative mass density differences

If perturbing the cytoskeleton, and the tensional balance between nucleus and cell boundary does not alter relative mass densities, what about changing chromatin compaction directly? After all, chromatin compaction is linked to transcriptional activity and controlled by epigenetic marks, which can be specifically altered using drugs.

We treated HeLa-FUCCI cells with trichostatin A (TSA) or anacardic acid (ANA) and measured their RI tomograms. TSA decondenses chromatin by inhibition of histone deacetylases (HDAC) (23), while ANA condenses chromatin by inhibition of histone acetyltransferases (HAT) (53). As shown in Fig. 5 *a*-*c* for the visual inspection and in Fig. 5 *d* and Supplementary Fig. 9 for the quantitative analysis, the TSA treatment decreased the RI of each cellular compartment while the ANA treatment induced the opposite effect.

**Figure 5.**
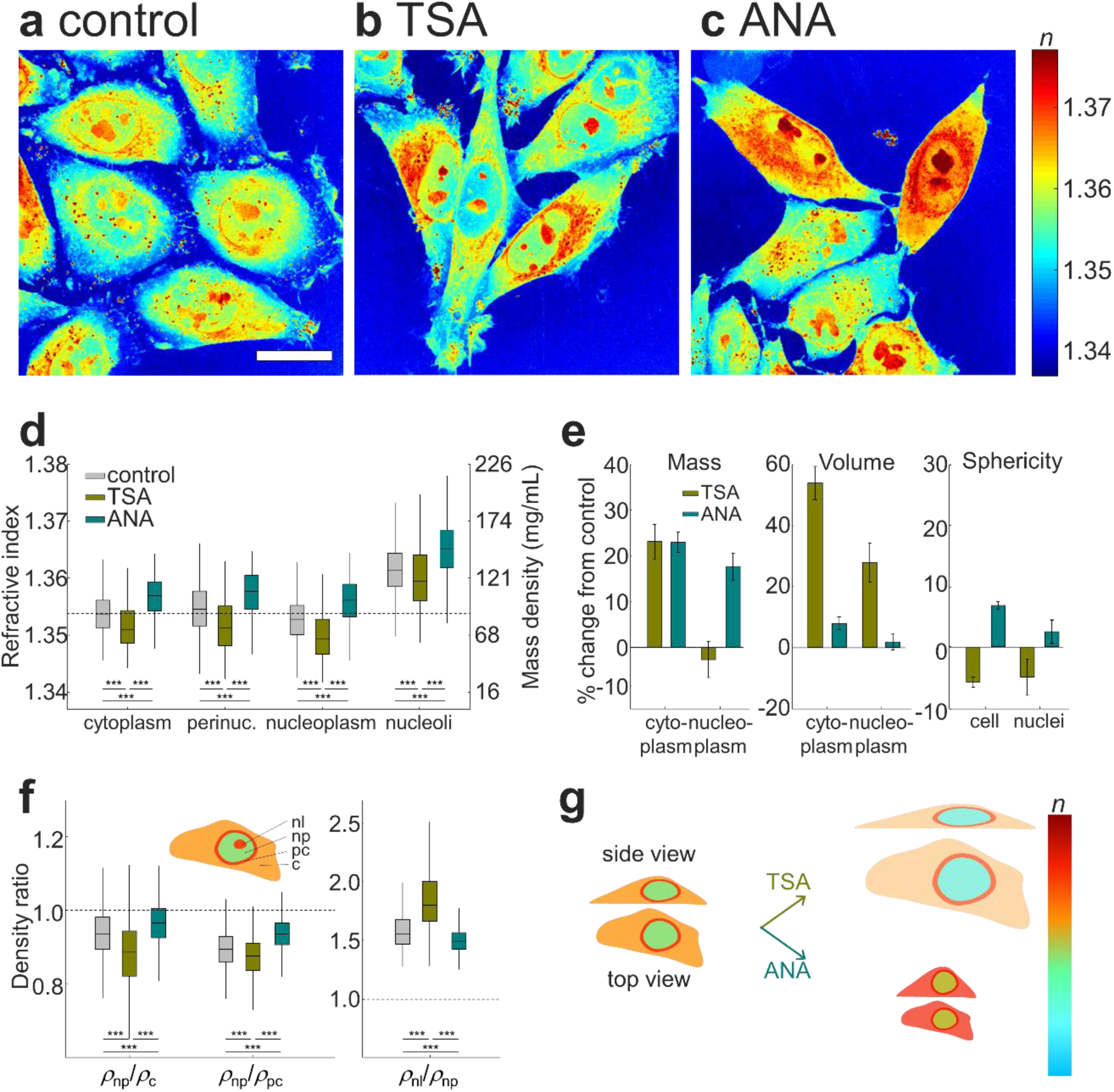
Changes in refractive index distribution of HeLa-FUCCI cells under chromatin condensation. (a-c) Maximum projection of RI tomograms of HeLa-FUCCI cells in (a) control, (b) TSA, and (c) ANA treatment. The scale bar is 20 µm. (d) Mean RI values of cytoplasm, perinuclear cytoplasm, nucleoplasm, and nucleoli in individual cells under chromatin condensation. The dashed line indicates the mean RI value of cytoplasm in control cells. (e) Relative changes of the mass, volume, and sphericity of the cytoplasm and nuclei under TSA and ANA treatment compared to control. (f) Mass density ratio between nucleoplasm and cytoplasm (*ρ*_np_/*ρ*_c_), nucleoplasm and perinuclear cytoplasm (*ρ*_np_/*ρ*_pc_), and nucleoli and nucleoplasm (*ρ*_nl_/*ρ*_np_) at the same phases of the cell cycle. The dashed lines indicate equal mass density of the compartments. The numbers of cells measured are *N* = 1,565, 437, and 924 for control, TSA, and ANA treatments, respectively. (g) Schematic indicating the effect of the TSA and ANA treatment on the volume, RI and shape of cells.

Quantitatively, the mean RI values of cytoplasm and nucleoplasm treated by TSA were lower than controls as 1.3516 ± 0.0004 and 1.3500 ± 0.0004 (1.3538 ± 0.0002 and 1.3528 ± 0.0002 for cytoplasm and nucleoplasm in controls). The decrease of RI and mass density of nucleoplasm in TSA-treated cells was mostly contributed from the significant swelling of nucleoplasm (28.0 ± 6.4%) while its dry mass exhibited subtle changes (–2.7 ± 3.9%) (Fig. 5 *e*). We also found that the dry mass of cytoplasm increased significantly by 23.2 ± 3.8% while the extensive volume increase of cytoplasm by 54.0 ± 5.5% lowered the RI value of cytoplasm. The cellular and nuclear sphericity also decreased by 5.9 ± 0.9% and 5.0 ± 3.0% under the TSA treatment, which could be the result of increased volume at constant cell height. These results indicate that increased histone acetylation increases nuclear volume (54, 55), and also suggest that the increase of euchromatin in nucleoplasm under the TSA condition enhances the protein synthesis in cytoplasm and is accompanied by an increase of cytoplasmic dry mass. This view is supported by previous findings reporting that TSA treatment increases mRNA expression by two-fold (55). In contrast, the mean RI values of cytoplasm and nucleoplasm under the ANA treatment were higher than in control cells as 1.3567 ± 0.0002 and 1.3560 ± 0.0002, respectively. The increases of cytoplasmic and nuclear RI under the ANA treatment are mainly due to increases in dry mass as 23.1 ± 2.2% and 17.7 ± 2.9%, respectively, while their volume increases were insignificant (8.0 ± 2.0% and 1.9 ± 2.6%) as shown in Fig. 5 *e*. It can be speculated that perturbing chromatin condensation not only affects the density of nucleoplasm, but also alters gene expression and protein synthesis. Moreover, HDAC and HAT can lead to the acetylation/deacetylation of non-histone proteins such as microtubules (56–58). Thus, the inhibition of those histone-modifying enzymes can modify the stability of microtubules for regulating cell volume, which may also indirectly affect the volume of cytoplasm and nucleoplasm.

Interestingly, despite the direct effects on chromatin condensation, nucleoplasm was still less dense than cytoplasm (Fig. 5 *f*). Under the TSA treatment, nucleoplasm was 11.8 ± 0.9% less dense than cytoplasm, which was more pronounced than in control cells (6.3 ± 0.4%). In contrast, nucleoplasm under the ANA treatment was 3.9 ± 0.9% less dense than cytoplasm. In addition, nucleoplasm under the TSA and ANA treatment was 12.7 ± 0.6% and 6.6 ± 0.3% less dense than perinuclear cytoplasm, respectively, which were equivalent to 10.0 ± 0.5 mg/ml and 7.0 ± 0.3 mg/ml of mass density difference across the nuclear membrane. Our finding is graphically summarized in Fig. 5 *g*.

## Discussion

In this study, we investigated the mass densities of nucleoplasm, cytoplasm, and nucleoli of adherent cells in various conditions including cell cycle progression and perturbations of cytoskeleton polymerization and chromatin condensation. We employed combined ODT and epi-fluorescence microscopy to measure 3D RI distributions of cells, from which the mass density, volume, and dry mass of nucleoplasm, nucleoli, and cytoplasm were quantitatively characterized. We found that nucleoplasm was less dense than cytoplasm in the majority of populations of adherent HeLa (82.6%) and RPE (76.5%) cells. This relative mass density difference was maintained during the cell cycle by scaling the volume of both compartments with the increase of dry mass of genetic and cellular materials. Moreover, this relative mass density difference was robust against drug-induced perturbations including depolymerization of actin and microtubules as well as chromatin condensation and decondensation.

The present ODT provides RI tomograms of individual cells and subcellular organelles with a spatial resolution of 121 nm (lateral) and 444 nm (axial), which are determined by the NAs of the objective lens and condenser lens (30). Spatial resolution is an important aspect of light microscopy techniques. At present, many techniques are published that push the boundary of spatial resolution way beyond what was thought possible because of the diffraction limit (down to nanometers) (59). However, all these techniques rely on fluorescence labels and cannot directly assess the physical properties of the sample. Also, ODT has better resolution than a typical brightfield or phase contrast microscope because the sample is illuminated from many different angles, effectively increasing the numerical aperture and thus spatial resolution (27, 30). Another measure of quality in any physical measurement is the precision of the measurement. In our ODT setup, the systematic errors arise from phase measurement, fast Fourier transform, and missing cone artifact for tomogram reconstruction. Altogether the precision of our RI measurement comes down to 4.15 × 10^−5^, which is sufficient to pick up the differences between the various regions of the cell characterized in the study.

The regulation of volume and dry mass in cell growth is essential for maintaining homeostasis in biological samples, and our finding confirms that the mass density of both nucleoplasm and cytoplasm are maintained during cell cycle progression. Whether this is caused by an active or passive mechanism is unclear at this point and remains to be determined. The result is in agreement with previous studies reporting that the buoyant density of nuclei is independent of the cell cycle (60, 61). At present we do not know what the functional importance of this particular mass density distribution is, but the mere fact that it is surprisingly stable against perturbations suggests that there is one. Moreover, we revealed that the relative mass density difference between nucleoplasm and nucleoli is also maintained during the cell cycle. This finding may imply that the chemical potential for nucleoli formation is maintained during the cell cycle, which governs formation of nucleoli by liquid-liquid phase separation (1). Interestingly, the averaged mass density of the nucleus including nucleoplasm and nucleoli is comparable to that of cytoplasm and perinuclear cytoplasm. Hence, the result may suggest that the redistribution of mass inside the nucleus is caused by the formation of nucleoli, thus reducing the mass density of nucleoplasm. This could be further investigated by measuring the mass density of nucleoplasm and nucleoli whose material properties can be modulated by an optogenetic gelation strategy (62). Recent studies have shown that the cell growth rate is size-dependent and exhibits exponential growth (63–65), and the size of nucleoli scales with that of cells while nuclei grow slower in the development of *C. elegans* embryo (66). We expect that time-resolved ODT measurements of nuclei and nucleoli during cell cycle progression can reveal the effect of the mass density of both compartments on the growth rate during the cell cycle. It will also be interesting to look specifically at the beginning of mitosis, when nucleoli dissolve, or at cells in developing embryos when there are no nucleoli yet.

We also discovered that nucleoplasm was still less dense than cytoplasm and perinuclear cytoplasm even when cytoskeleton polymerization or chromatin condensation was perturbed by drug treatments. In addition, we revealed that the mass density difference between nucleoplasm and cytoplasm exhibited subtle changes on the order of few mg/ml against the drug-induced perturbations. Perhaps these slight alterations of mass density difference result from mechanical force imbalances on the nuclear membrane in response to perturbations of cytoskeletal polymerization and chromatin condensation. On nuclear membrane, an osmotic pressure gradient generated by the mass density difference between nucleoplasm and perinuclear cytoplasm is mechanically balanced with pressure exerted by the cytoskeleton. The cytoskeletal pressure consists of a compressive (inward) component contributed by microtubules and a tensile (outward) pressure by the actomyosin cytoskeleton. There is also an additional elastic component from the mechanical stiffness of the nuclear envelope (21, 50–52, 67, 68). The osmotic pressure in solution, *Π*, is determined by the Van’t Hoff equation as *Π* = *ρRT*/*M* where *ρ* is the mass density of solutes, *R* is the gas constant (8.314 J·mol^-1^·K^-1^), *T* is the temperature of the solution, and *M* is the molecular weight of solutes. Hence, we can imagine that the slight mass density changes in two compartments alter their volume fraction (*ρ*/*M*) difference across the nuclear membrane under various perturbations, which can modify the osmotic pressure difference on the nuclear envelope, adding to the pressure exerted by the cytoskeleton. One way to decode the underlying biophysical principle of how and why nucleoplasm maintains a lower mass density than the surrounding cytoplasm could be to manipulate the osmotic pressure in the nucleoplasm and cytoplasm experimentally (69, 70) and determining the changes of the mass density of the compartments.

The nucleus is a very interesting organelle from a materials perspective. It combines a highly packed, topologically constrained and highly charged polymeric material in its bulk with an internally and externally interconnected envelope, and is subject to many active processes (71). Some of its internal structure also derives from liquid-liquid phase separation. Thus, it does not have to be surprising that density is not linearly related to stiffness. A simple gedankenexperiment might help to illustrate the point: a crosslinked and a non-crosslinked polymer gel have the same density, but clearly different elasticity and viscosity. The nucleus has even been shown to behave like an auxetic material with a negative Poisson ratio (72). So, our finding of a low mass density is not in conflict with the fact that it generally has a high elastic modulus. Moreover, the nucleus mechanically interacts with the surrounding cytoplasm and exterior environment via the cytoskeleton. The effects of the nuclear and cytoplasmic mass densities on their mechanical properties, and *vice versa*, is still unexplored, primarily due to the lack of practical techniques to measure local mechanical properties inside cells. There are emerging non-invasive optical microscopic techniques to probe the mechanical properties of biological samples including Brillouin microscopy (52, 73, 74), fluorescence lifetime imaging (FLIM) (75, 76), and time-lapse quantitative phase microscopy (77). In the future, combining ODT with such other microscopic modalities can reveal the interaction between the mass density and mechanical properties of nucleus and cytoplasm.

To conclude, we have combined ODT and epi-fluorescence microscopy to investigate the mass density of nucleoplasm, cytoplasm, and nucleoli of adherent cells. We revealed that nucleoplasm is less dense than cytoplasm while nucleoli have significantly higher mass density than nucleoplasm. Moreover, the relative mass density difference between the compartments was robustly maintained during cell cycle progression and upon perturbations of cytoskeletal integrity and chromatin condensation. Biological cells are increasingly understood as physical objects amenable to physical modelling, and the mass density of a particular region or organelle in a cell might appeared as a fundamental physical parameter in such physical models. Thus, providing quantitative values for such physical parameters has a value in itself, as it can then be incorporated in the physical models. Although the physiological relevance of the phenomenon of lower mass density of the nucleoplasm versus the cytoplasm needs further investigation, our findings based on the quantitative characterization may have implications for understanding cell homeostasis during the cell cycle, the mechanical interaction between nucleus and cytoplasm via the cytoskeleton, and nuclear volume regulation. Further studies are also required to identify whether the robustness of the relative mass densities is due to active regulatory mechanisms by the cell, or whether they are an emerging result of basic physico-chemical properties of the cell.

## Author contributions

K.K. realized the combined optical setup for optical diffraction tomography and epi-fluorescence microscopy and conducted measurements. K.K. and J.G. designed the experiments, interpreted the results, and wrote the manuscript.

## Acknowledgements

We thank Vasily Zaburdaev, Simone Reber, Simon Alberti, Francis Stewart, Hendrik Dietz, Ada and Don Olins, Paul Janmey, Gheorghe Cojoc, Raimund Schlüßler, Paul Müller, Anna Taubenberger, Joan-Carles Escolano, Shada Abuhattum, Marta Urbanska, Abin Biswas, and Timon Beck for helpful discussions and technical support. HeLa-FUCCI and RPE-FUCCI cell lines were kindly provided by the lab of Frank Buchholz (TU Dresden). We also thank Ina Poser in the lab of Anthony Hyman (Max Planck Institute of Molecular Cell Biology and Genetics, Dresden) for providing the stable HeLa cell line with nucleoli stained with GFP. The authors acknowledge financial support from the Volkswagen Foundation (research grant 92847 to KK and JG) and the Alexander von Humboldt Stiftung (Alexander von Humboldt Professorship to JG). The authors declare no conflict of interest.

---

**Supplementary Figure 1.**
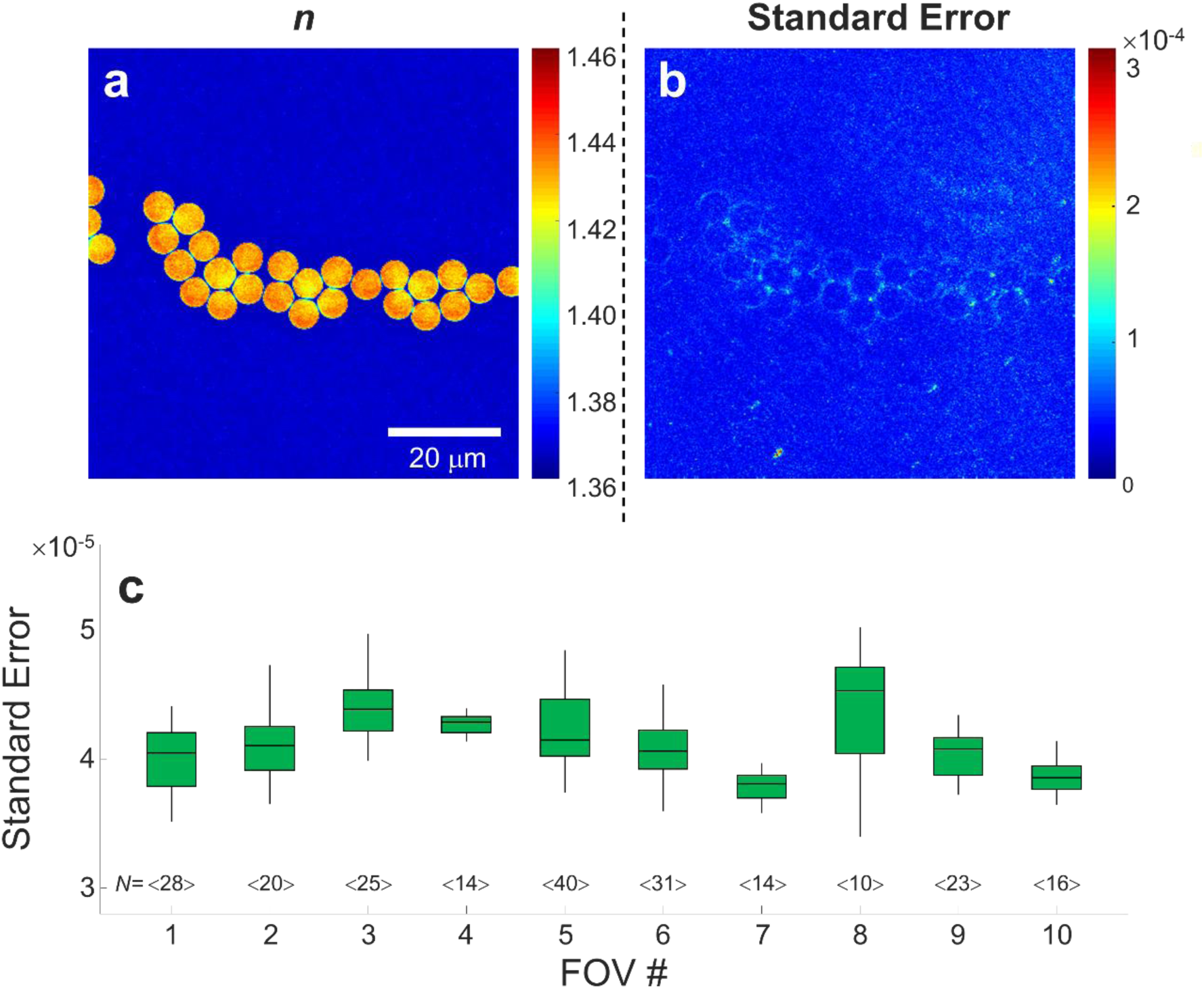
Precision of ODT measurements. The cross-sectional slice of (a) an RI tomogram of silica beads immersed in 0.7 M sucrose solution and (b) the standard error of a time series of 10 RI tomograms of the same silica beads. (c) The averaged standard error of a time series of 10 RI tomograms within individual silica bead in each field-of-view (FOV). The number of silica beads in an individual FOV is indicated as *N*. The averaged standard error of RI is calculated as 4.15 × 10^−5^.

**Supplementary Figure 2.**
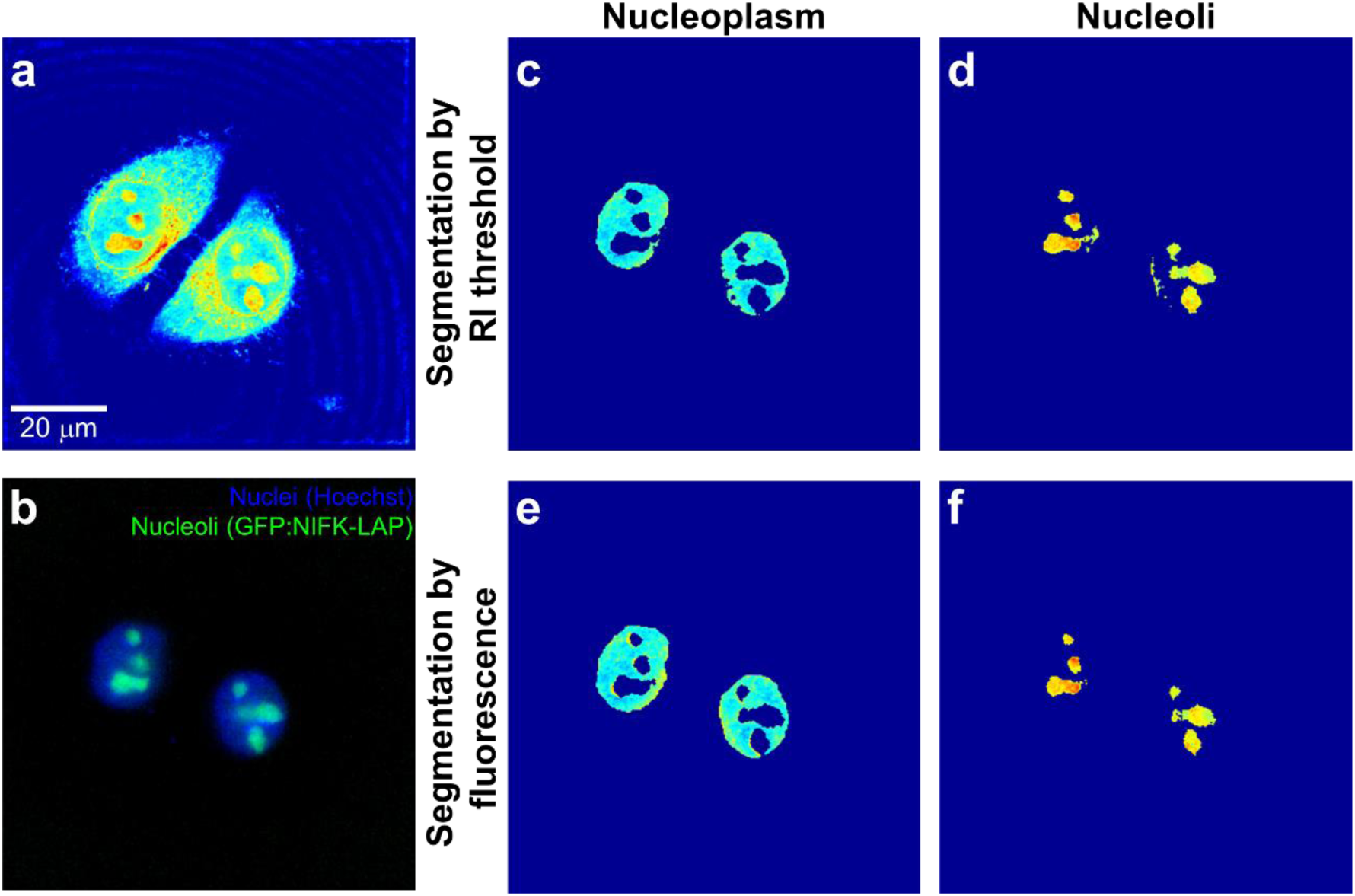
Confirmation of the segmentation of nucleoli based on the RI threshold. (a-b) The measured (a) RI tomogram and (b) epi-fluorescence image of HeLa cells which nuclei and nucleoli are stained with Hoechst and GFP, respectively. (c-d) The cross-sectional slices of the RI tomogram of (c) nucleoplasm and (d) nucleoli segmented by the present method based on the RI threshold. (e-f) The cross-sectional slices of the RI tomogram of (e) nucleoplasm and (f) nucleoli segmented by the fluorescence intensity.

**Supplementary Figure 3.**
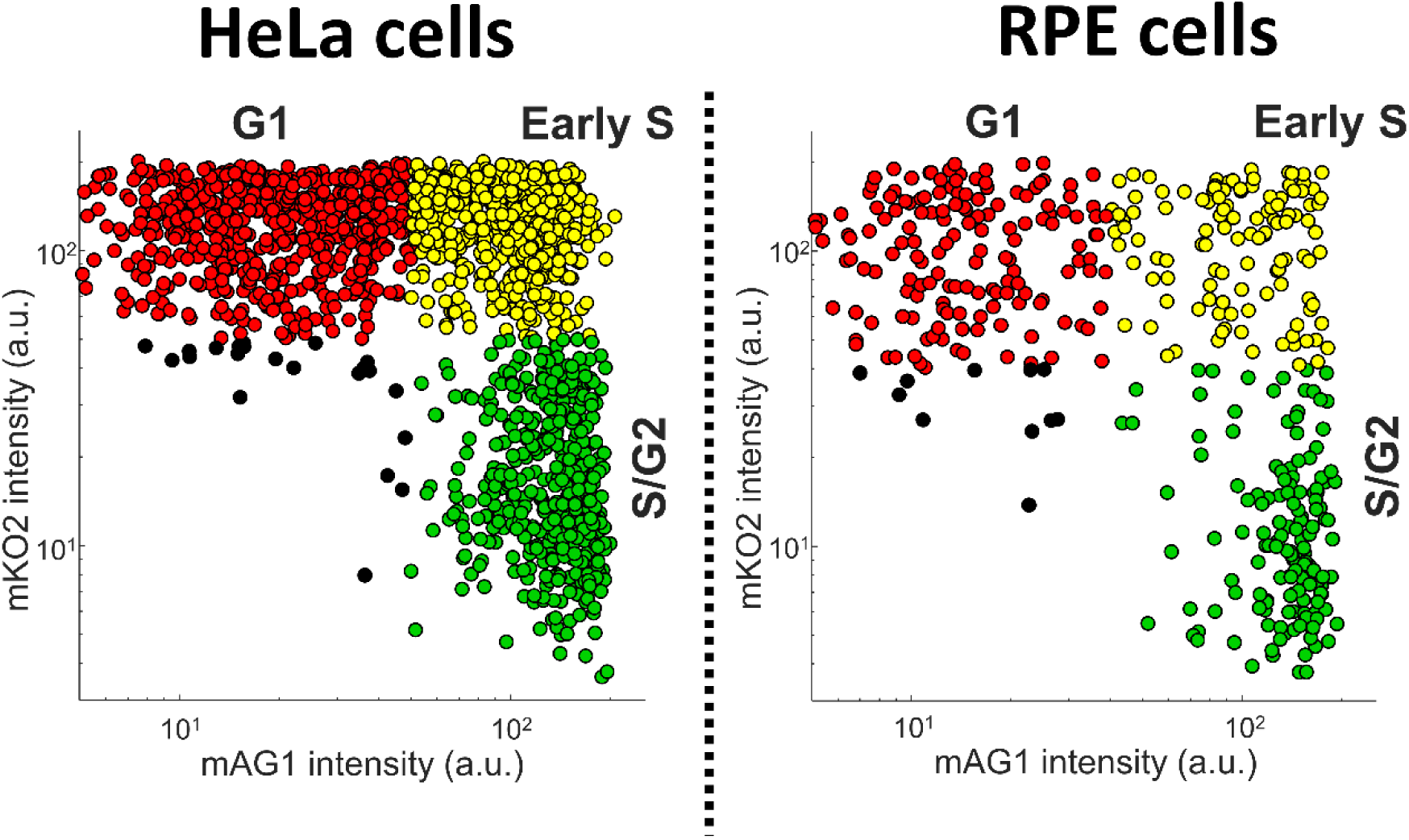
Determination of cell cycle phases of individual cells by FUCCI cell-cycle marker. The two-channel epi-fluorescence microscopy measured the intensity of fluorescence signals in FUCCI-stained nucleus in cells (mAG1 and mKO2), and the cell cycle phase of cells was determined in the two-color scatter plot as G1 (high mKO2 signals), early S (high mKO2 and mAG1 signals), and S/G2 (high mAG1 signals) phases. The black dots are determined as either cells in mitosis or cells not expressing fluorescence signals, and discarded in statistical analysis.

**Supplementary Figure 4.**
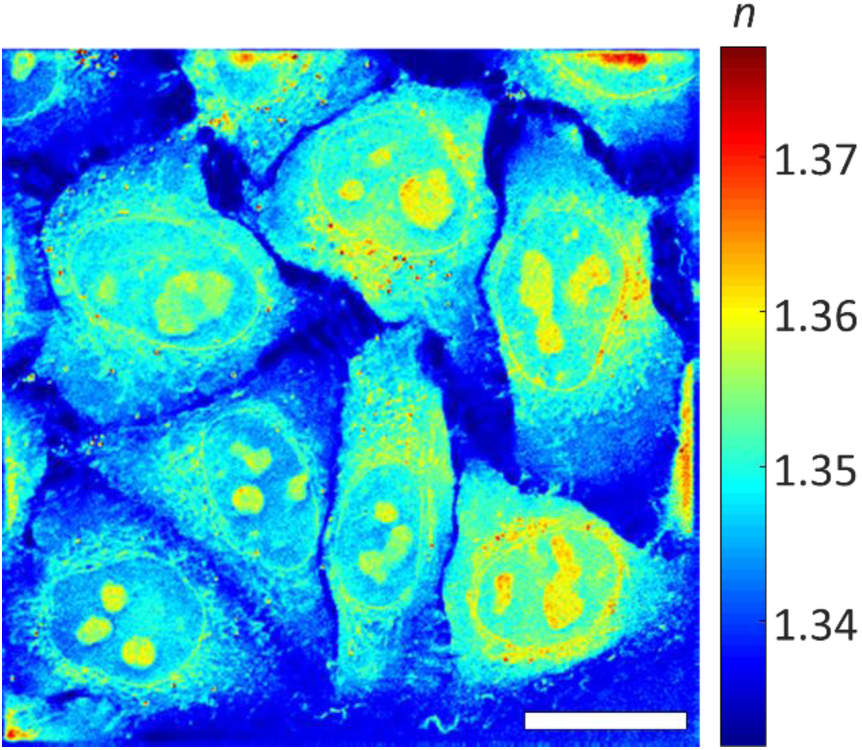
Refractive index (RI) and mass density distributions in non-labeled HeLa cells. The maximum projection of an RI tomogram of non-labeled HeLa cells in the direction proportional to the substrate. The scale bar is 20 µm.

**Supplementary Figure 5.**
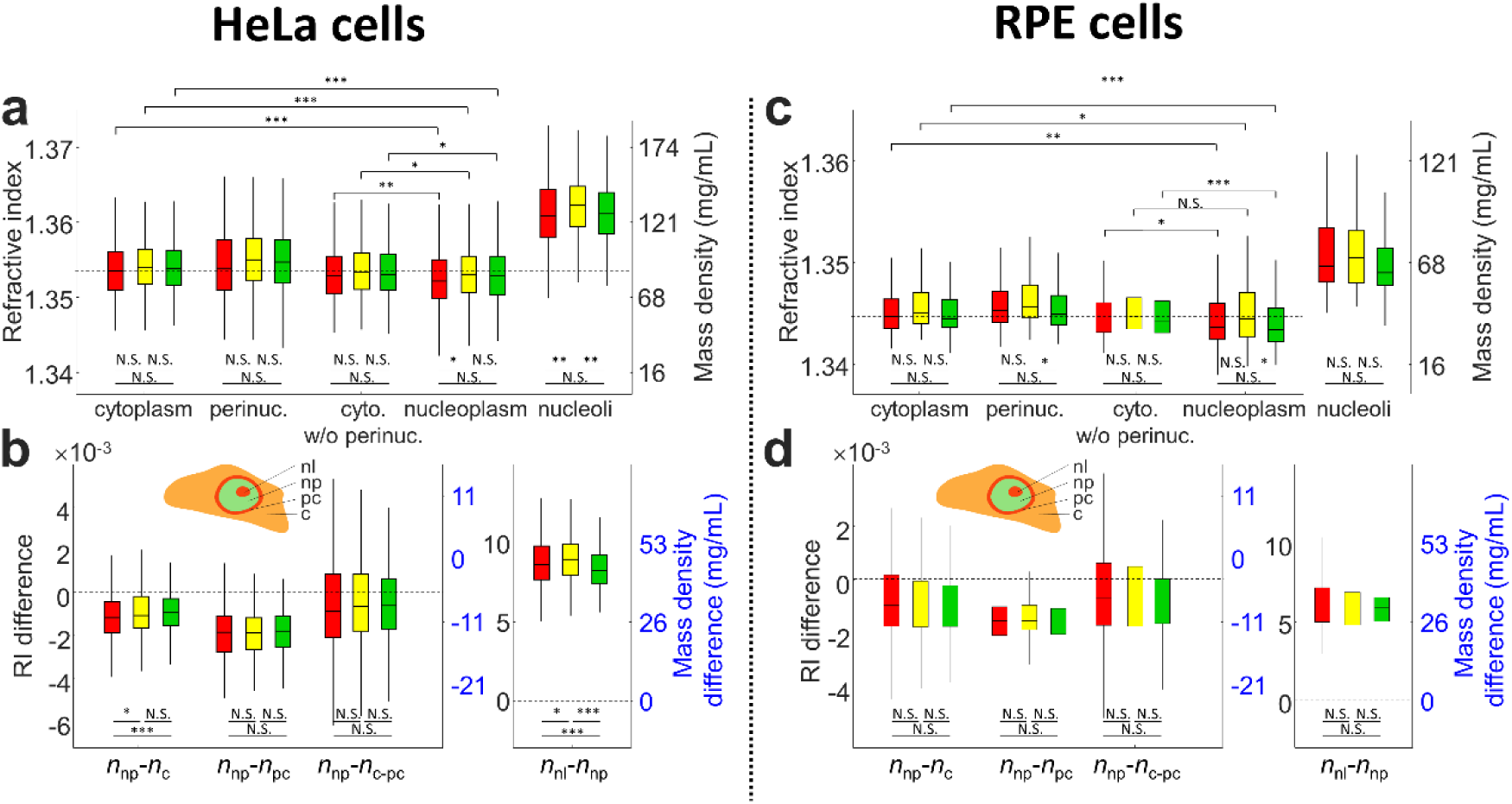
Refractive index (RI) and mass density distributions in HeLa-FUCCI (a-b) and RPE-FUCCI (c-d) cells as function of the cell cycle. (a, c) The mean RI values of cytoplasm, perinuclear cytoplasm, cytoplasm excluding perinuclear region, nucleoplasm, and nucleoli. The dashed line indicates the mean RI value of cytoplasm at the G1 phase in the cell cycle. (b, d) The difference of the mean RI values between nucleoplasm and cytoplasm (*n*_np_-*n*_c_), nucleoplasm and perinuclear cytoplasm (*n*_np_-*n*_pc_), nucleoplasm and cytoplasm excluding perinuclear region (*n*_np_-*n*_c-pc_), and nucleoli and nucleoplasm (*n*_nl_-*n*_np_). The dashed lines indicate the equal mass density of the compartments. The numbers of cells measured are *N* = 557, 505, and 483 for HeLa cells, and *N* = 122, 92, and 128 for RPE cells in the G1, early S, and S/G2 phases, respectively.

**Supplementary Figure 6.**
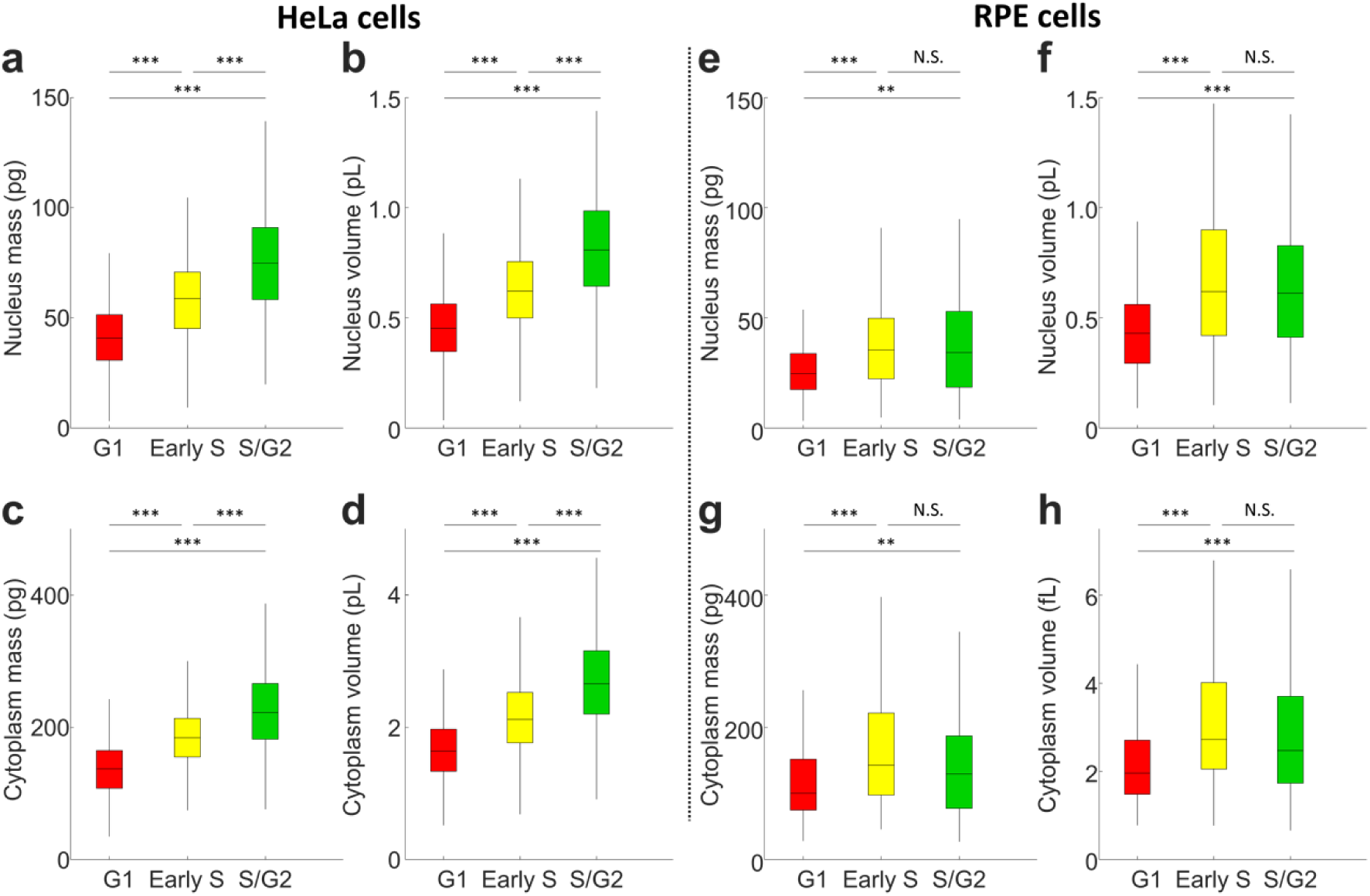
Mass and volume distribution of HeLa-FUCCI (a-d) and RPE-FUCCI (e-h) cells as function of the cell cycle. (a, e) The mass and (b, f) volume of nucleus, and (c, g) the mass and (d, h) volume of cytoplasm in HeLa-FUCCI and RPE-FUCCI cells, respectively. The numbers of cells measured are *N* = 557, 505, and 483 for HeLa cells, and *N* = 122, 92, and 128 for RPE cells in the G1, early S, and S/G2 phases, respectively.

**Supplementary Figure 7.**
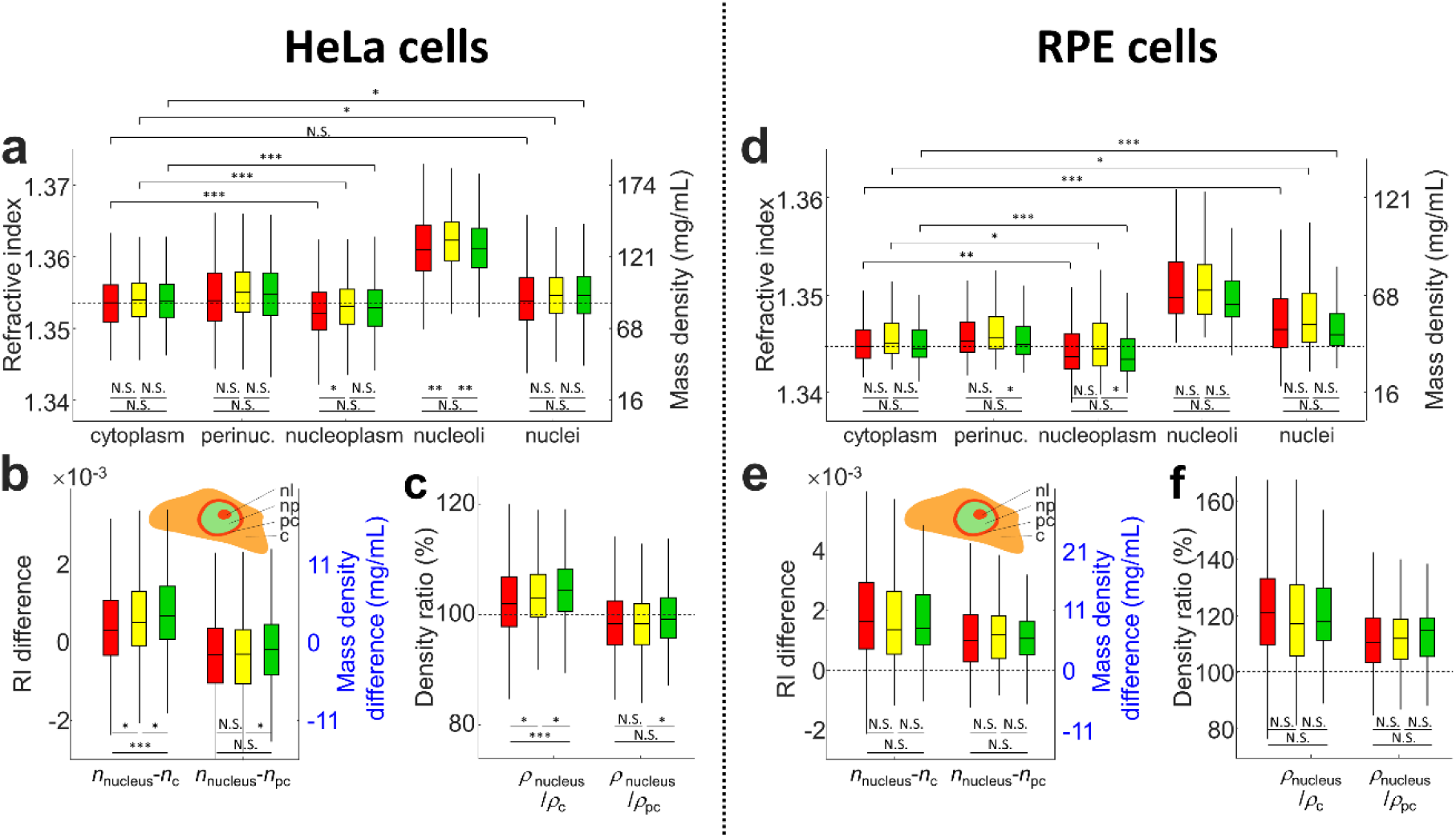
Refractive index (RI) and mass density distributions in HeLa-FUCCI (*a*-*c*) and RPE-FUCCI (*d*-*f*) cells as function of the cell cycle. (*a, c*) The mean RI values of cytoplasm, perinuclear cytoplasm, nucleoplasm, nucleoli, and nuclei including nucleoplasm and nucleoli. The dashed line indicates the mean RI value of cytoplasm at the G1 phase in the cell cycle. (*b, d*) The difference of the mean RI values between nucleus and cytoplasm (*n*_np_-*n*_c_) and nucleus and perinuclear cytoplasm (*n*_np_-*n*_pc_). The dashed lines indicate the equal mass density of the compartments. (*c, f*) The ratio of the mass density between nucleus and cytoplasm (*ρ*_nucleus_/*ρ*_c_) and nucleus and perinuclear cytoplasm (*ρ*_nucleus_/*ρ*_pc_). The dashed lines indicate equal mass density between compartments. The numbers of cells measured are *N* = 557, 505, and 483 for HeLa cells, and *N* = 122, 92, and 128 for RPE cells in the G1, early S, and S/G2 phases, respectively.

**Supplementary Figure 8.**
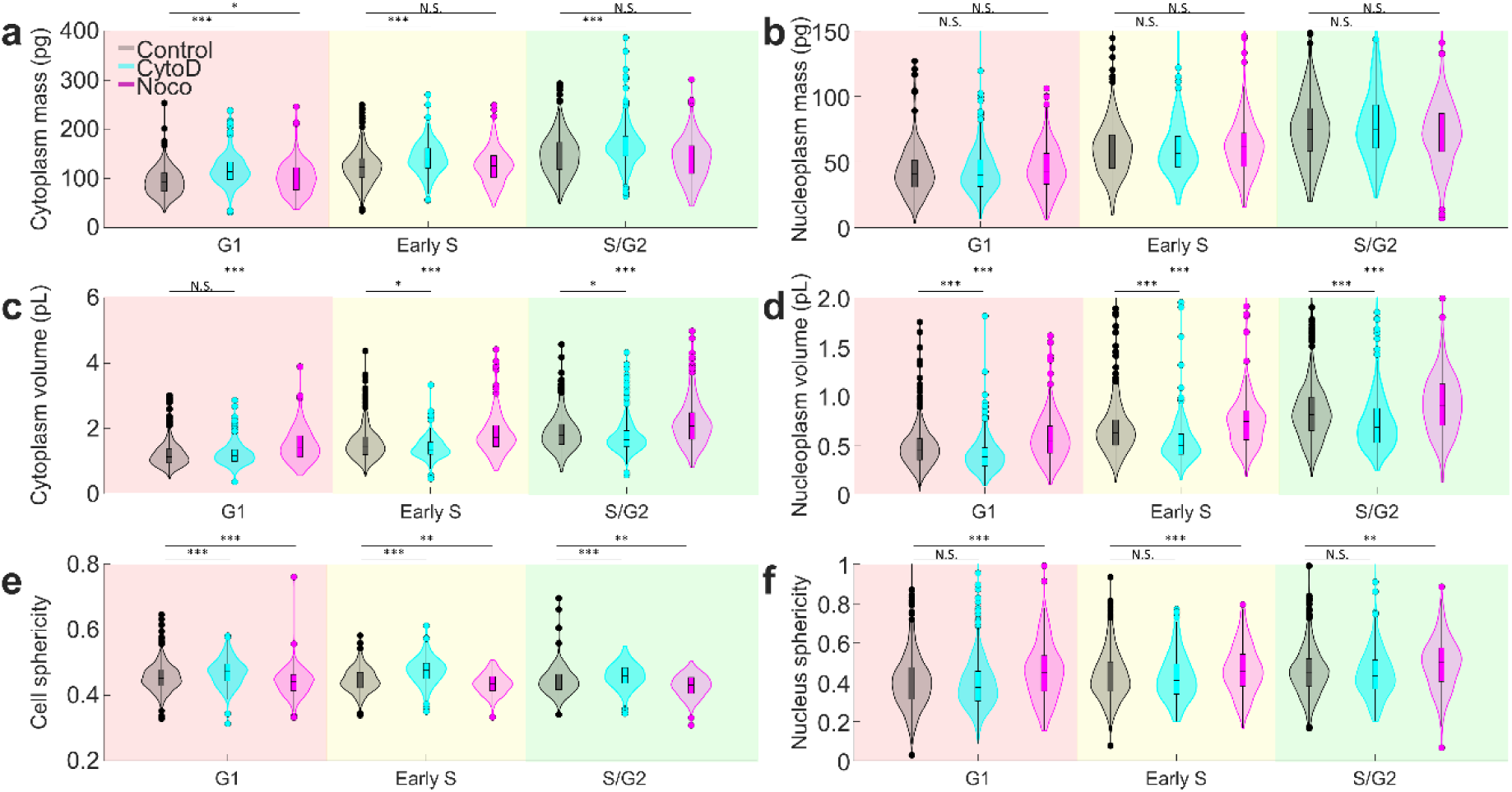
Changes in mass, volume, and sphericity of HeLa-FUCCI cells under cytoskeletal perturbation. (a-d) The mass and volume of nucleus and cytoplasm in HeLa-FUCCI cells in control (gray), cytoD (cyan) and noco (magenta) treatment at various cell cycle phases. (e-f) The sphericity of whole cell (e) and nucleus at various cell cycle phases. The cell cycle phases are indicated as shaded region in red (G1), yellow (Early S) and green (S/G2). The numbers of cells measured are *N* = 1,565, 973, and 717 for control, cytoD, and nocodazole treatments, respectively.

**Supplementary Figure 9.**
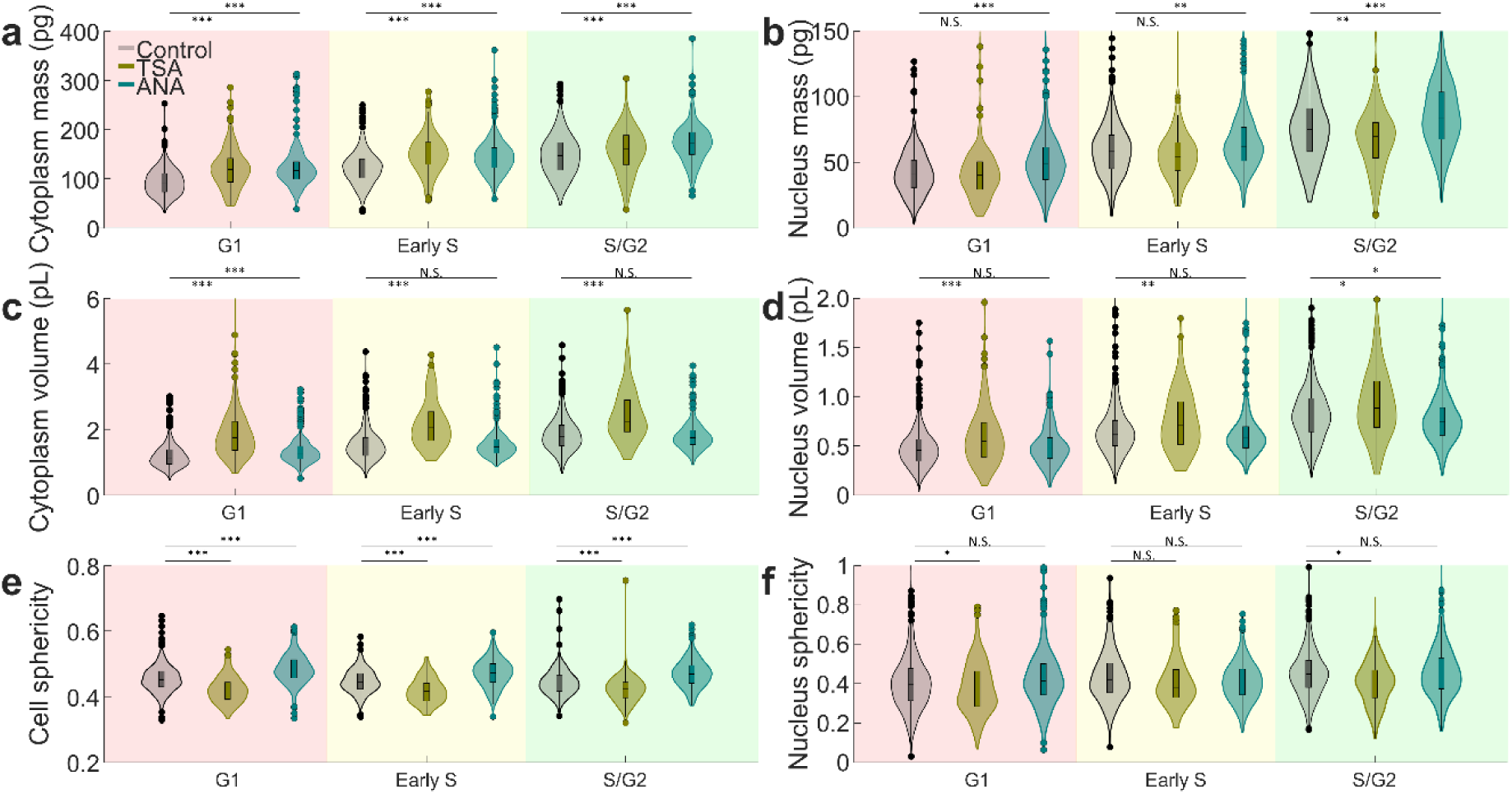
Changes in mass, volume, and sphericity of HeLa-FUCCI cells under chromosome condensation. (a-d) The mass and volume of nucleus and cytoplasm in HeLa-FUCCI cells in control (gray), TSA (dark yellow) and ANA (dark green) treatment at various cell cycle phases. (e-f) The sphericity of whole cell (e) and nucleus at various cell cycle phases. The cell cycle phases are indicated as shaded region in red (G1), yellow (Early S) and green (S/G2). The numbers of cells measured are *N* = 1,565, 437, and 924 for control, TSA, and ANA treatments, respectively.

